# The self-organized learning of noisy environmental stimuli requires distinct phases of plasticity

**DOI:** 10.1101/612341

**Authors:** Steffen Krüppel, Christian Tetzlaff

## Abstract

Along sensory pathways, representations of environmental stimuli become increasingly sparse and expanded. If additionally the feed-forward synaptic weights are structured according to the inherent organization of stimuli, the increase in sparseness and expansion leads to a reduction of sensory noise. However, it is unknown how the synapses in the brain form the required structure, especially given the omnipresent noise of environmental stimuli. Here, we employ a combination of synaptic plasticity and intrinsic plasticity - adapting the excitability of each neuron individually - and present stimuli with an inherent organization to a feed-forward network. We observe that intrinsic plasticity maintains the sparseness of the neural code and thereby enables synaptic plasticity to learn the organization of stimuli in low-noise environments. Nevertheless, even high levels of noise can be handled after a subsequent phase of readaptation of the neuronal excitabilities by intrinsic plasticity. Interestingly, during this phase the synaptic structure has to be maintained. These results demonstrate that learning in the presence of noise requires adaptation of the synaptic structure but also of the neuronal properties in two distinct phases of learning: an *encoding phase*, during which the inherent organization of the environmental stimuli is learned, followed by a *readaptation phase* to readapt the neuronal system according to the current level of noise. The necessity of these distinct phases of learning suggests a new role for synaptic consolidation.

## Introduction

Learning to distinguish between different stimuli despite high levels of noise is an important ability of living beings to ensure survival. However, the underlying neuronal and synaptic processes of this ability are largely unknown.

The brain is responsible for controlling movements of an agent’s body in response to the perceived stimulus. For instance, the agent should run away from a predator or run after the prey. To do so, the agent needs to be able to reliably classify the perceived stimulus despite its natural variability (e.g. different individuals of the same predator species) or noise (e.g. impaired vision by obstacles). In general, the sensory processing systems of the brain map the stimulus representation onto subsequent brain areas yielding successive representations which are increasingly sparse in activity and expansive in the number of neurons. If the feed-forward synaptic weights realizing this mapping are structured according to the inherent organization of the stimuli (e.g. lion versus pig), the increased sparseness and expansion lead to a significant reduction of noise and therefore to a reliable classification [1]. However, it remains unclear how the synapses form the required structure despite noise during learning. Furthermore, how can the system reliably adapt to varying levels of noise (e.g. being in a silent forest compared to near a loud stream)?

In the mouse olfactory system, for instance, 1800 glomeruli receiving signals from olfactory sensory neurons project to millions of pyramidal neurons in the piriform cortex yielding an expansion of the stimulus representation [2, 3]. Activity of the glomeruli is relatively dense with 10%-30% of glomeruli responding to a given natural odor [4], while in the piriform cortex activity drops to 3%-15% indicating an increase in sparseness [5, 6]. A similar picture can be observed in the drosophila olfactory system. Here, 50 glomeruli project to about 2500 Kenyon cells in the mushroom body [7, 8]. While about 59% of projection neurons respond to a given odor, only 6% of Kenyon cells do [9]. Similar ratios have been observed in the locust olfactory system [10]. In the cat visual system, the primary visual cortex has 25 times as many outputs than it receives inputs from the LGN [11]. In addition, V1-responses to natural visual stimuli are significantly sparser than in the LGN [12, 13]. Both principles of increased expansion and sparseness of stimulus representations apply to other sensory processing systems as well [14–16].

The functional roles of increased sparseness as well as expansion have already been proposed in the Marr-Albus theory of the cerebellum [17, 18]. Here, different representations are thought to evoke different movement responses although the activity patterns overlap. The Marr-Albus theory demonstrates that through expansion and the sparse activity of granule cells, the overlapping patterns are mapped onto non-overlapping patterns that can easily be classified. A recent theoretical study has focused on sparse and expansive feed-forward networks in sensory processing systems [1]. Here, small variations in activity patterns are caused by internal neuronal noise, input noise, or changes in insignificant properties of the stimuli. For reliable stimulus classification, these slightly varying activity patterns belonging to the same underlying stimulus should evoke the same response in a second layer (or brain area) of a sparse and expansive feed-forward network. Surprisingly, although the network is sparse and expansive, random synaptic weights increase both noise and overlap of activity patterns in the second layer. On the other hand, the same network with synaptic weights structured according to the organization of stimuli reduces the noise and overlap of activity patterns simplifying subsequent classification. How a network is able to learn the organization of stimuli, shape its synaptic structure according to this organization, and do so even in the presence of noise is so far unknown.

The generally accepted hypothesis of learning is that it is realized by changes of synaptic weights by the process of (long-term) synaptic plasticity [19, 20]. Synaptic weights are strengthened or weakened depending on the activity of the pre- and postsynaptic neurons [21–23]. Hebbian plasticity describes the process of increasing a synaptic weight if the activity of the two connected neurons is correlated [19]. Several theoretical studies indicate that Hebbian plasticity alone would lead to divergent synaptic and neuronal dynamics thus requiring homeostatic synaptic plasticity [24, 25] to counterbalance and stabilize the dynamics [26–30]. In addition, neurons adapt their excitability by the processes of intrinsic plasticity [31, 32]. Intrinsic plasticity regulates the excitability of a given neuron so as to maintain a desired average activity [33–36]. Although several studies indicate that intrinsic plasticity, for instance, optimizes the input-output relation of a neuron [31] or the encoding of information in firing rates [37], its specific role in learning, especially in sensory systems, remains unclear.

In the present study, we show that intrinsic plasticity regulates the neuronal activities such that the synaptic weights can learn the organization of stimuli even in the presence of low levels of noise. Interestingly, after learning, the system is able to adapt itself according to changes in the level of noise it is exposed to - even if these levels are high. To do so, intrinsic plasticity has to readapt the excitability of the neurons while the synaptic weights have to be maintained indicating the need of a two-phase learning protocol.

In the following, first, we present the basics of our theoretical model and methods and demonstrate the ability of a feed-forward network with static random or static structured synaptic weights, respectively, to distinguish between noisy versions of 1000 different stimuli (similar to [1]). Then, we introduce the synaptic and intrinsic plasticity rules considered in this study. We train the plastic feed-forward network during an *encoding phase* without noise and test its performance afterwards by presenting stimuli of different noise levels. Intriguingly, the self-organized dynamics of synaptic and intrinsic plasticity yield a performance and network structure similar to the static network initialized with structured synaptic weights. Further analyses indicate that the performance of the plastic network to classify noisy stimuli greatly depends on the neuronal excitability, especially for high levels of noise. Hence, after learning without noise, we changed the noise level in order to test the performance but let intrinsic plasticity readapt the excitability of the neurons. This *readaptation phase* significantly increases the performance of the network. Note, however, if synaptic plasticity is present during this second phase, the increase in performance is impeded by a prolonged and severe performance decrease. In the next step, we show that in the encoding phase with both intrinsic and synaptic plasticity the network can also learn from noisy stimuli if the level of noise is low. Again, high levels of noise impede learning and classification performance. However, after the subsequent readaptation phase the network initially trained with low-noise stimuli is also able to classify highly noisy stimuli.

## Results

### Model setup and classification performance

The main question of this study concerns how sparse and expansive neural systems, such as diverse sensory processing areas, learn the inherent organization of stimuli enabling a reduction of noise? To tackle this question, similar to a previous study [1], we consider a neural network that consists of two layers of rate-based neurons with the first layer being linked to the second layer via all-to-all feed-forward synaptic connections (Fig 1A). The first layer, called stimulus layer, is significantly smaller (*N*_*S*_ = 1000 neurons) than the second one, called cortical layer, (*N*_*C*_ = 10000 neurons) and the activity patterns of the stimulus layer serve as stimuli or inputs to the cortical layer. These stimulus patterns are constructed of firing rates 𝒮_*i*_ = {0, 1} of the stimulus neurons *i* = {1, …, *N*_*S*_} with “0” representing a silent neuron and “1” a maximally active one. Neurons belonging to the cortical layer posses a membrane potential *u*_*j*_ (*j* = {1, …, *N*_*C*_}) modeled by a leaky integrator receiving the inputs from the stimulus layer. The membrane potential of a cortical neuron is transformed into a firing rate 𝒞_*j*_ using a sigmoidal transfer function. Similar to the stimulus neurons, we consider the minimal and maximal firing rates ℱ^min^ = 0 and ℱ^max^ = 1. Note that the point of inflection of the sigmoidal transfer function *ε*_*j*_, also called cortical firing threshold, is neuron-specific.

**Fig 1.**
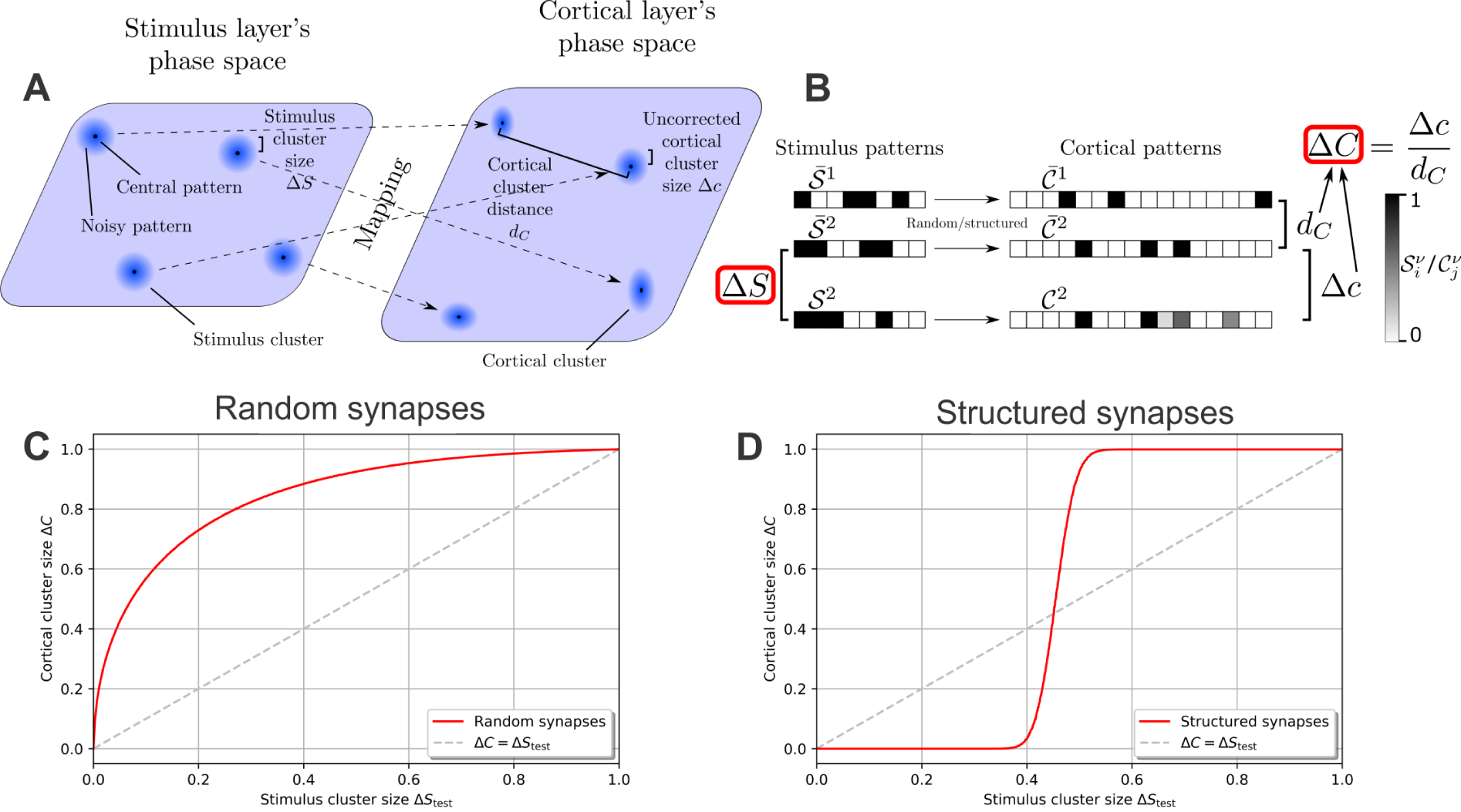
Network model and mathematical approach to quantifying the noise reduction by an expansive, sparse network. **(A)** The feed-forward network consists of two layers of rate-based neurons with the stimulus layer projecting stimuli onto the cortical layer via all-to-all feed-forward synaptic connections. Stimuli are organized in *P* = 1000 clusters, with each cluster *ν* consisting of a characteristic central pattern 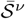 (black dots) and noisy patterns 𝒮^*ν*^ (blue halos around dots). The size of the stimulus clusters Δ*S* corresponds to the level of noise and is indicated schematically by the size of the blue halos. Stimulus clusters are mapped by the synaptic connections to cortical clusters containing central cortical patterns 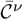 and noisy cortical patterns 𝒞^*ν*^. **(B)** Illustration of different patterns and measures used in this study. The activity of each neuron (box) is indicated by its gray scale (left: stimulus layer; right: cortical layer). The central pattern of each stimulus cluster (underlying stimulus) evokes a specific central pattern in the cortical layer. Noisy versions of a central stimulus pattern (here 𝒮^2^) activate different cortical patterns with their average distance Δ*c* from the original pattern depending on the structure of the feed-forward synaptic weights. **(C)** Random synaptic weights increase the cluster size for all stimulus cluster sizes, i.e. Δ*C* > Δ*S*_test_; the noise in the stimuli is thus amplified by the network. **(D)** Synapses that are structured in relation to the organization of the underlying stimuli (stimulus central patterns 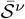)decrease the size of clusters, i.e. the noise, up to a medium noise level (Δ*S*_test_ ≈ 0.45). (C,D) Dashed line indicates Δ*C* = Δ*S*_test_.

The different activity patterns of the stimulus layer are organized into *P* = 1000 stimulus clusters. Each stimulus cluster *ν* = {1, …, *P*} consists of one characteristic activity pattern, called central stimulus pattern 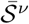, which represents the underlying stimulus (e.g. a lion; black dots in the stimulus layer’s phase space in Fig 1A). A central stimulus pattern is constructed by assigning each stimulus neuron *i* a firing rate 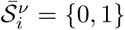 with equal probability (see Fig 1B for schematic examples). In addition, a stimulus cluster contains all noisy versions 𝒮^*ν*^ of the underlying stimulus (e.g. a lion behind a tree or a rock; indicated by blue halos in Fig 1A) generated by randomly flipping firing rates 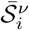 of the cluster’s central stimulus pattern from “1” to “0” or vice versa with probability Δ*S/*2 (Fig 1B). Δ*S* thus reflects the average noise level of all noisy stimulus patterns as well as the stimulus cluster’s size in the stimulus layer’s phase space. If Δ*S* = 0, the cluster is only a single point in the stimulus layer’s phase space (the central stimulus pattern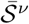) and thus it is noise-free. The maximum value of the stimulus cluster size Δ*S* = 1 represents a cluster that is distributed evenly across the entire phase space. Here, the noise is so strong that no information remains. The stimulus cluster size Δ*S* can be retrieved by the normalized Hemming distance between patterns:

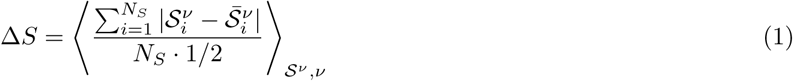

with the brackets denoting the average over all noisy stimulus patterns 𝒮^*ν*^ of all stimulus clusters *ν*.

Every activity pattern of the stimulus layer elicits an activity pattern in the cortical layer, such that stimulus clusters are mapped to cortical clusters (dashed arrows in Fig 1A). Similar to the stimulus cluster, each cortical cluster consists of one central pattern 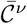 (evoked by the noise-free stimulus 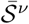) and noisy patterns 𝒞^*ν*^ (evoked by the noisy stimuli 𝒮^*ν*^). Due to the complex mapping of the stimulus patterns onto the cortical layer via the feed-forward synaptic weights, it is not clear how the level of noise is affected by this mapping. Therefore, we estimate the noise in the cortical layer in analogy to Eq (1):

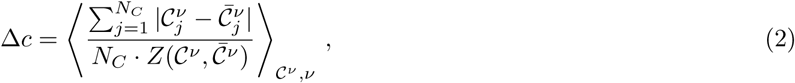

where 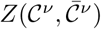 is a normalization factor (see Methods for more details). As different stimulus clusters are mapped by the same feed-forward weights onto cortical clusters, random correlations between the cortical clusters could be induced. To account for these correlations we calculate the average distance between clusters by

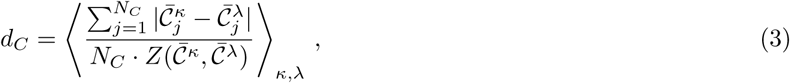

and correct Eq 2 using this cortical cluster distance (Eq 3) analogous to a signal-to-noise/noise-to-signal ratio to obtain the cortical cluster size

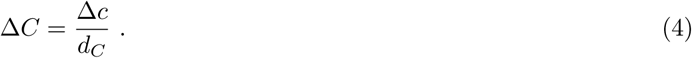

Therefore, if each pattern 𝒮^*ν*^ of the same stimulus cluster *ν* is mapped onto a different (random) pattern 𝒞^*ν*^ in the cortical layer, Δ*C* = 1 and the cluster is distributed evenly over the entire cortical layer’s phase space. If each pattern of a stimulus cluster is mapped onto the same pattern of the cortical cluster (the central pattern 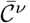), Δ*C* = 0.

In summary, both the stimulus cluster size Δ*S* as well as the cortical cluster size Δ*C* are measures for the amount of random fluctuations of different activity patterns belonging to the same underlying stimulus. As such, a network tasked with reducing these random fluctuations should decrease the cluster size, i.e. Δ*C <* Δ*S*.

### Static networks

Central to the performance in reducing the cluster size or noise are the feed-forward synaptic weights *ω*_*ji*_ between neurons. In the following, we predefine the synaptic weights and test the performance of the network for different levels of noise Δ*S*_test_ while keeping the synaptic weights fixed. For each noise level Δ*S*_test_, we create noisy stimulus patterns for all clusters and use them to evaluate the average noise Δ*C* in the cortical layer. By doing so, we obtain a performance curve Δ*C* (Δ*S*_test_) of the network. If the weights are initialized randomly (here drawn from a Gaussian distribution 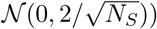, the cortical cluster size Δ*C* is always larger than the stimulus cluster size Δ*S*_test_, as the performance curve is above the identity line (Δ*C* = Δ*S*_test_; dashed line in Fig 1C) for all values of Δ*S*_test_. In other words, the noise of the stimuli (Δ*S*_test_) is amplified by the network by increasing the variations between different cortical patterns of the same underlying stimulus (Δ*C* > Δ*S*_test_). Note that this amplification of noise is present although the network is expansive and sparse [1].

This picture changes when the weights are structured according to the organization of environmental stimuli. To portray such a structure, we initialize the synaptic weights according to [1, 38]:

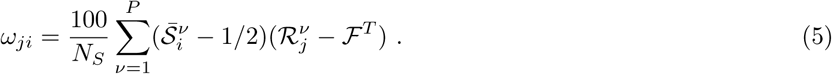

This results in a mapping of the central stimulus patterns 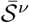 to randomly generated, ℱ^*T*^ -sparse cortical patterns ℛ^*ν*^ (ℱ^*T*^ = 0.001). Interestingly, this mapping yields a reduction of noise for up to medium levels (Δ*S*_test_ ≲ 0.45) such that the cortical cluster size Δ*C* is smaller than the stimulus cluster size Δ*S*_test_ (Fig 1D). In other words, as already shown in a previous study [1], a structured network reduces small fluctuations of representations of the same underlying stimulus. Note that in the random as well as the structured network the cortical neurons’ average activity at Δ*S*_test_ = 0 is about 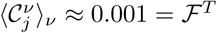 (see Methods). We chose this value as it results in all cortical neurons of the structured network firing in response to exactly one central stimulus pattern, and remaining silent in response to all others (as ℱ^*T*^ *P* = 1), which simplifies the qualitative analysis of the results.

These results show that expansive and sparse networks reduce the noise of stimuli if the synaptic weights from the stimulus to the cortical layer are structured according to the underlying organization of stimuli (here according to the central stimulus patterns 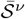). However, by using Eq 5 the synaptic weights are set on the required values - they are not learned given the environmental stimuli.

### Plastic network

As demonstrated above, a network with random synaptic weights increases the level of noise, while a structured network decreases it (Fig 1C,D). How can a network develop this structure in a self-organized manner given only the environmental stimuli? To investigate this question, we initialized a network with the same random synaptic weights as above, i.e. Gaussian distributed *ω*_*ji*_, and let the system evolve over time using plasticity mechanisms that adapt the synaptic weights and neuronal excitabilities. These plasticity mechanisms are assumed to depend on local quantities only and thus on the directly accessible neuronal activities and synaptic weights [27, 39]. Given this assumption, the environmental stimuli influence the dynamics of the plasticity mechanisms as the stimulus patterns determine the activities of the neurons. We consider two plasticity processes: Synaptic weights are controlled by Hebbian correlation learning and an exponential decay term (for weight stabilization),

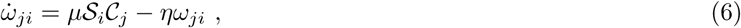

while a faster intrinsic plasticity mechanism regulates the firing thresholds *ε*_*j*_ of the cortical neurons so as to achieve the target firing rate ℱ^*T*^ = 0.001:

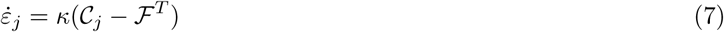

with the parameters *μ, η, κ* determining the time scales of the mechanisms.

Training is carried out in repeated learning steps or trials. In each learning step *L*, we present all central stimulus patterns 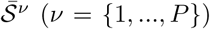 to the network once, ensuring there is no chronological information (see Methods for details). This corresponds to a stimulus cluster size Δ*S*_learn_ = 0 or noise-free learning. At different stages of learning (that is, after different numbers of learning steps) we test the performance of the network for different levels of noise Δ*S*_test_ like has been done for the static networks.

As learning progresses (Fig 2A), the performance curve develops from the random network’s (red line), which amplifies stimulus noise, into one similar to the structured network’s performance curve (blue compared to magenta line). As will be shown in the following, the plasticity mechanisms (Eqs 6 and 7) enable the network to encode the organization of the stimuli (existence of different clusters) in a self-organized manner, with most of the performance gained in the first *L* = 60000 learning steps. During learning, the synaptic weights evolve from the initial Gaussian distribution into a bimodal distribution with peaks at about 0.33 and 0 (see Fig 2B for an example). The emergence of the bimodal weight distribution and its link to the network performance can be explained as follows: Due to the random initialization of the synaptic weights each central stimulus pattern leads to a different membrane potential in a given cortical neuron (see, e.g., Fig 2D) such that all *P* stimuli together yield a random distribution of evoked membrane potentials. As the target firing rate is chosen such that each neuron ideally responds to only one central stimulus pattern (as ℱ^*T*^ *P* = 1), intrinsic plasticity adapts the firing threshold *ε*_*j*_ of a neuron such that one of the evoked membrane potentials leads to a distinctly above-average firing rate. Consequently, synapses connecting stimulus neurons, which are active at the corresponding stimulus pattern, with the considered cortical neuron are generally strengthened the most by Hebbian synaptic plasticity. These synapses will likely form the upper peak of the synaptic weight distribution (Fig 2B). Meanwhile, all other synaptic weights are dominated by the synaptic weight decay (second term in Eq 6) and will later form the lower peak of the distribution at zero. As the continued differentiation of the synaptic weights increases the evoked membrane potential of the most influential central stimulus pattern, these two processes of synaptic and neuronal adaptation drive each other. Interestingly, the resulting synaptic weights are correlated to the structured synapses (Fig 2C) initialized using Eq 5 (here the cortical patterns ℛ^*ν*^ of Eq 5 were generated using the central cortical patterns 𝒞^*ν*^ of the plastic network at the corresponding learning step *L*; see Methods for further details). Note that the cortical firing thresholds *ε*_*j*_ of the plastic network become correlated to the values of the static, structured one as well (Fig 2E).

**Fig 2.**
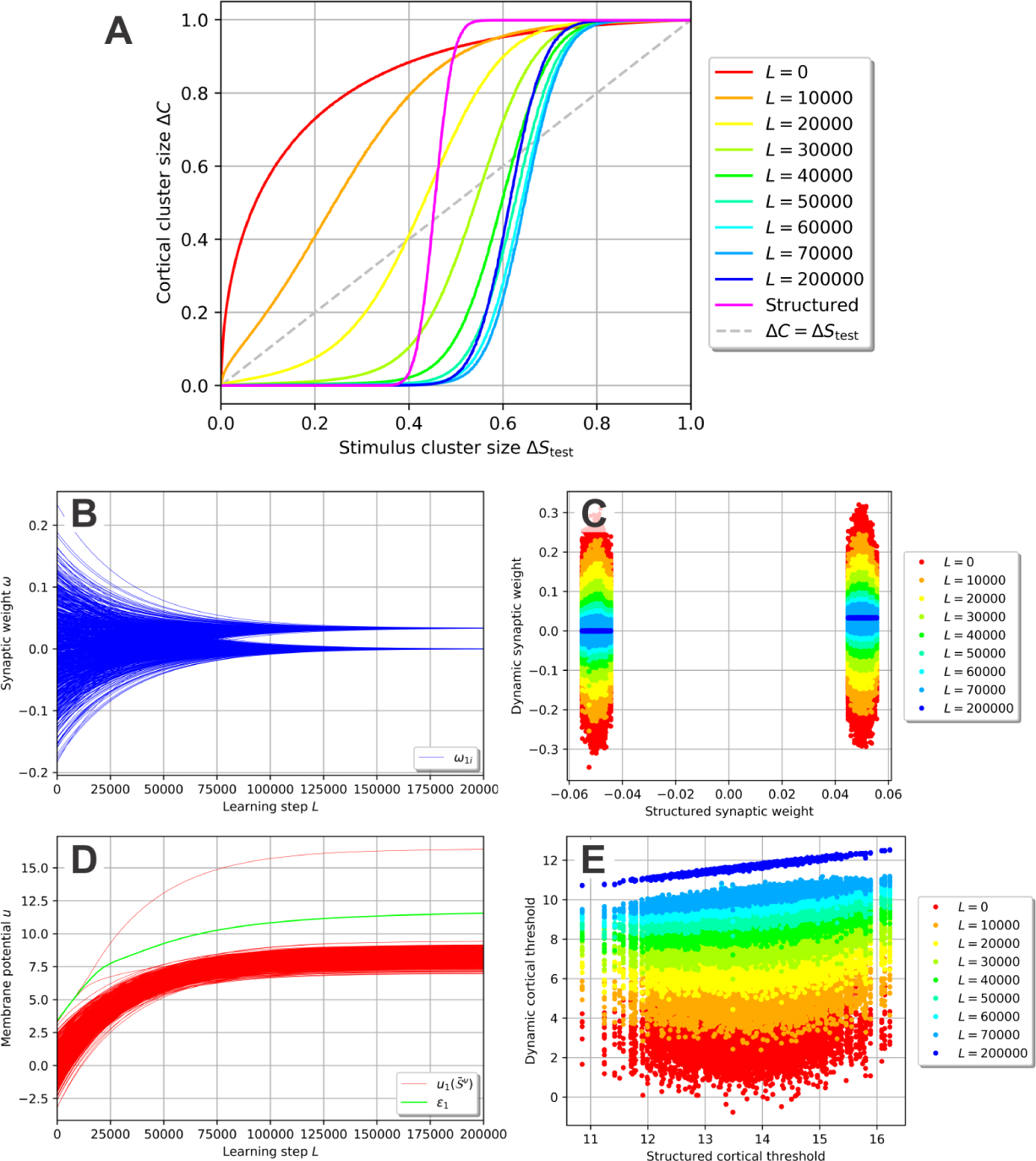
Self-organization of the synaptic and neuronal structure via synaptic and intrinsic plasticity in a noise-free environment. **(A)** By repeatedly presenting one stimulus pattern *𝒮*^*ν*^ per cluster per learning step *L* using a stimulus cluster size Δ*S*_learn_ = 0 (i.e. presenting the central stimulus patterns 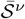), the network’s performance develops from the noise-amplification of a random network (red, equal to Fig 1C) to a performance significantly decreasing the level of noise for Δ*S*_test_ up to about 0.6 (blue). **(B,C)** During learning, the synaptic weights develop into a bimodal distribution (B; only the weights connecting to neuron 1 are shown) which is correlated to the distribution of the static, structured network (C). **(D)** For each cortical neuron (here shown for neuron 1), the firing threshold (green) increases such that only one central stimulus pattern can evoke a membrane potential larger than the threshold. **(E)** Similar to the synaptic weights (C), the firing thresholds tend to become correlated to the ones of the static, structured network.

In summary, synaptic and intrinsic plasticity interact and adapt the neuronal network such that, in a noise-free environment, it learns to encode the organization of the stimuli in a way comparable to a static, pre-structured network.

### The functional role of the cortical firing thresholds

While being structurally similar, the performance of the trained, plastic network (Fig 2A, blue) appears significantly better than the performance of the static, structured network (magenta). This fact is not self-explanatory, since both the synaptic weights as well as the cortical firing thresholds are strongly correlated between both networks (Fig 2C,E). However, a closer look at the cortical firing thresholds and their link to the performance of the network reveals the cause of this difference as discussed in the following.

In the trained network (*L* = 200000), as mentioned before, each cortical neuron should fire in response to the central stimulus pattern 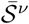 of exactly one cluster and stay silent otherwise. Exemplary, we will focus on cortical neuron *j* = 1 which fires in response to the central stimulus pattern 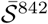 of cluster *ν* = 842 and remains silent in response to all other central stimulus patterns. In general, two types of errors can occur.

#### *False negatives* (a stimulus of cluster 842 is presented and cortical neuron 1 falsely does not fire)

Noisy patterns of cluster 842 elicit a distribution of membrane potentials in cortical neuron 1 (Fig 3A), which depends on the stimulus cluster size Δ*S*_test_, i.e. the level of noise. All noisy stimulus patterns 𝒮^842^ which evoke a membrane potential in neuron 1 that is higher than the neuron’s firing threshold *ε*_1_ result in a strong activation of neuron 1. The neuron therefore classifies these 𝒮^842^ correctly as belonging to cluster 842. However, noisy patterns 𝒮^842^ evoking a lower membrane potential than *ε*_1_ do not elicit strong activation of cortical neuron 1. These noisy patterns are falsely classified as not belonging to cluster 842 and correspond to false negatives.

**Fig 3.**
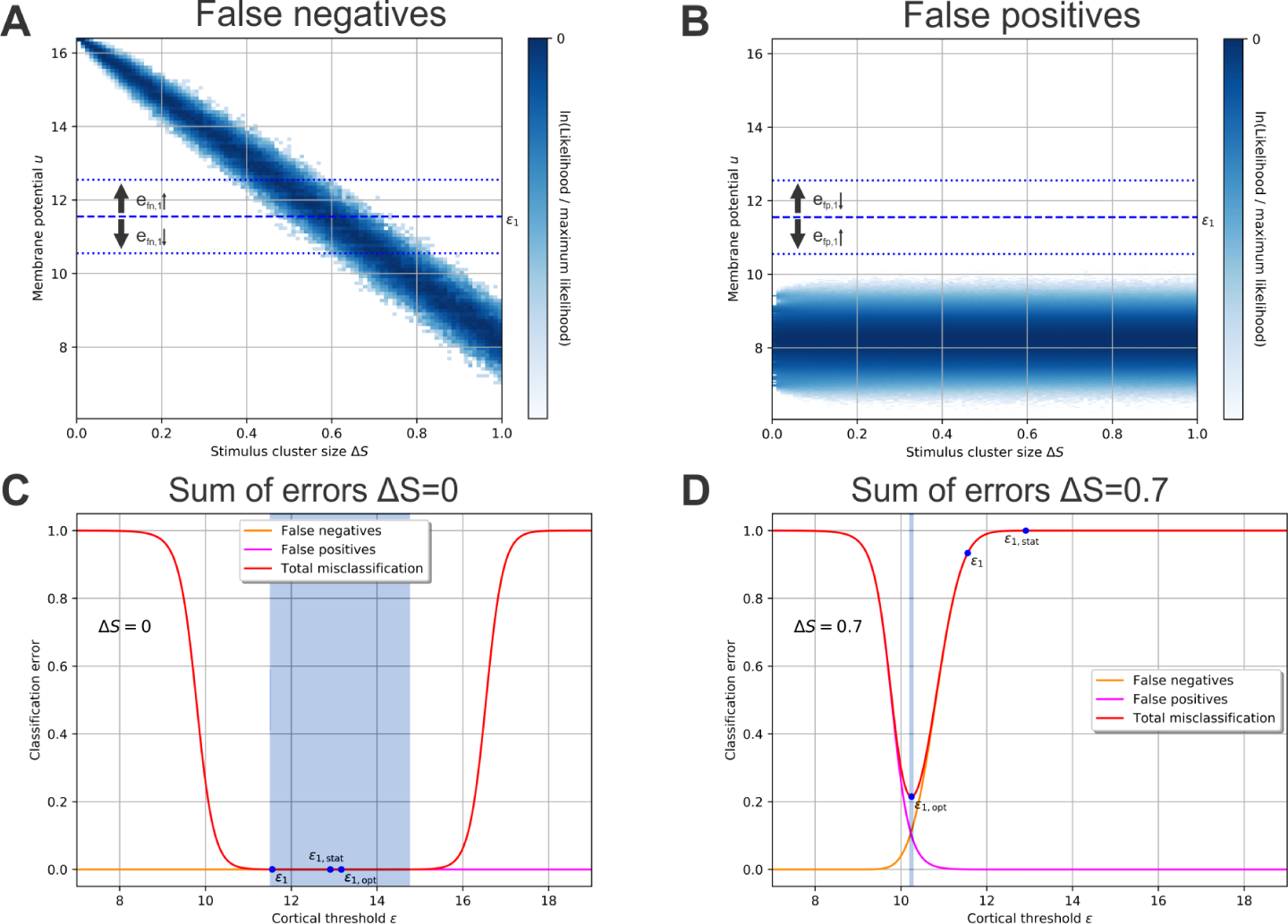
The Classification performance of each neuron depends on its firing threshold. In a single cortical neuron (here neuron *j* = 1), multiple noisy stimulus patterns of the same stimulus cluster elicit a distribution of membrane potentials. Two distinct distributions can be identified: **(A)** The distribution of membrane potentials evoked by noisy stimulus patterns belonging to the cluster whose central pattern elicits firing in the given cortical neuron (here cluster *ν* = 842). For any Δ*S*_test_, all stimuli yielding a membrane potential that is below the neuron’s firing threshold (dashed line; *ε*_1_) do not elicit a strong neuronal response representing false negatives. The distribution significantly depends on the level of noise Δ*S*_test_. **(B)** The membrane potential distribution in response to noisy stimulus patterns of the clusters the neuron is not tuned to (*ν* ≠ 842). Here, all stimuli yielding a membrane potential above the firing threshold are false positives. **(C)** Δ*S*_test_ = 0: A higher firing threshold *ε* leads to more false negatives (orange) but fewer false positives (magenta) and vice versa for a lower threshold. The sum of errors (red) is negligible in a large regime (blue area: gradient is less than 0.001). **(D)** Δ*S*_test_ = 0.7: With higher levels of stimulus noise, the total error and the classification performance depend critically on the firing threshold. (C,D) *ε*_1,opt_: optimal value of the firing threshold for the given level of noise Δ*S*_test_ yielding the lowest total error; *ε*_1_: value of the firing threshold after learning with noise-free stimuli (Δ*S*_test_; Fig 2); *ε*_1,stat_: firing threshold in the static network (Fig 1D).

#### *False positives* (a stimulus of a cluster *ν* ≠ 842 is presented and cortical neuron 1 falsely fires)

Similar to the analysis of false negatives, the analysis of false positives can be done with clusters whose central patterns should not elicit activity in cortical neuron 1. The distribution of membrane potentials evoked by noisy patterns of these clusters does not significantly depend on the stimulus cluster size Δ*S*_test_ (Fig 3B). Noisy stimulus patterns 𝒮^*ν*^ (*ν* ≠ 842) are classified correctly as not part of cluster 842 if neuron 1’s membrane potential is lower than its firing threshold *ε*_1_. All noisy patterns evoking a higher membrane potential falsely lead to a firing of cortical neuron 1. They correspond to false positives.

Both false positives and false negatives depend on the firing threshold *ε*_*j*_ of a neuron *j*. For all values of Δ*S*_test_, a lower firing threshold would generally lead to less false negatives (*e*_fn,*j*_; Fig 3A) but simultaneously to more false positives (*e*_fp,*j*_; Fig 3B) and vice versa for a higher firing threshold. Consequently, there is a trade-off between false negatives and false positives with their sum being related to the network’s performance or cortical cluster size (see Methods):

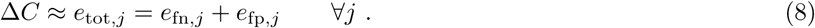

The performance of the network or the total error *e*_tot,*j*_ thus depends on a cortical neuron’s firing threshold in a nonlinear manner. Given noise-free stimuli (Δ*S*_test_ = 0), in a large regime of different values for the firing threshold cortical neuron 1 makes almost no classification error (red line in Fig 3C; between vertical blue lines the gradient is less than 0.001). For a higher noise level (e.g. Δ*S*_test_ = 0.7, Fig 3D), there is no such extended regime of low-error threshold values. Instead, small variations of the firing threshold can drastically change the classification performance, since the membrane potential response distributions overlap at these noise levels (Fig 3A,B).

During training without noise (Δ*S*_learn_ = 0), the neuronal firing threshold *ε*_1_ rose to the lower bound of the low-error regime of Δ*S*_test_ = 0 (blue area; Fig 3C). In the static network, however, firing thresholds *ε*_1,stat_ were placed at the center of the highest and second highest membrane potentials in response to central stimulus patterns leading to much higher values. Therefore, if the network performance is tested for small stimulus clusters (low noise Δ*S*_test_; Fig 3C), the static and the plastic network have a similar total error and classification performance. For larger stimulus clusters (high noise levels Δ*S*_test_; Fig 3D), on the other hand, the higher firing thresholds of the static network lead to considerably more misclassification and consequently to a higher cortical cluster size Δ*C*. Consequently, the fact that the relation between the threshold and its classification error *e*_*j*,tot_ depends on the noise Δ*S*_test_ provides an explanation for the large performance differences between the static structured and the plastic network (Fig 2A).

This example (Fig 3) demonstrates that the value of the neuron-specific threshold *ε*_*j*,opt_ optimizing a neuron’s classification performance depends on the stimulus cluster size Δ*S*_test_ or current level of noise (dotted lines in Fig 4A for neuron 1 in blue and neuron 2 in green). The firing thresholds after training (solid lines in Fig 4A), however, are independent of Δ*S*_test_, as they are determined by the noise present during training (Δ*S*_learn_ = 0). For Δ*S*_test_ ≲ 0.5 these thresholds are within the regime of low total error (shaded areas indicate the low-error regime for each neuron marked by blue lines in Fig 3C,D) yielding a high classification performance of the network. However, for Δ*S*_test_ ≳ 0.5 the thresholds resulting from training without noise (Δ*S*_learn_ = 0) start to deviate significantly from the optimal thresholds leading to a decreasing classification performance (Fig 2A and Fig 4C solid lines for total error of individual neurons). Interestingly, the deviation from the optimal threshold is accompanied by a decrease of the average activity level (solid lines; Fig 4B), while the optimal thresholds would keep the cortical activity close to the target activity ℱ^*T*^ = 0.001 (dotted lines; for Δ*S*_test_ ≳ 0.85 the total error is high and nearly independent of the threshold, see S1 Fig). We thus expect that after initial learning intrinsic plasticity could readapt the neuronal firing thresholds according to the present level of noise such that the target activity is maintained and the thresholds approximate the optimal threshold values.

**Fig 4.**
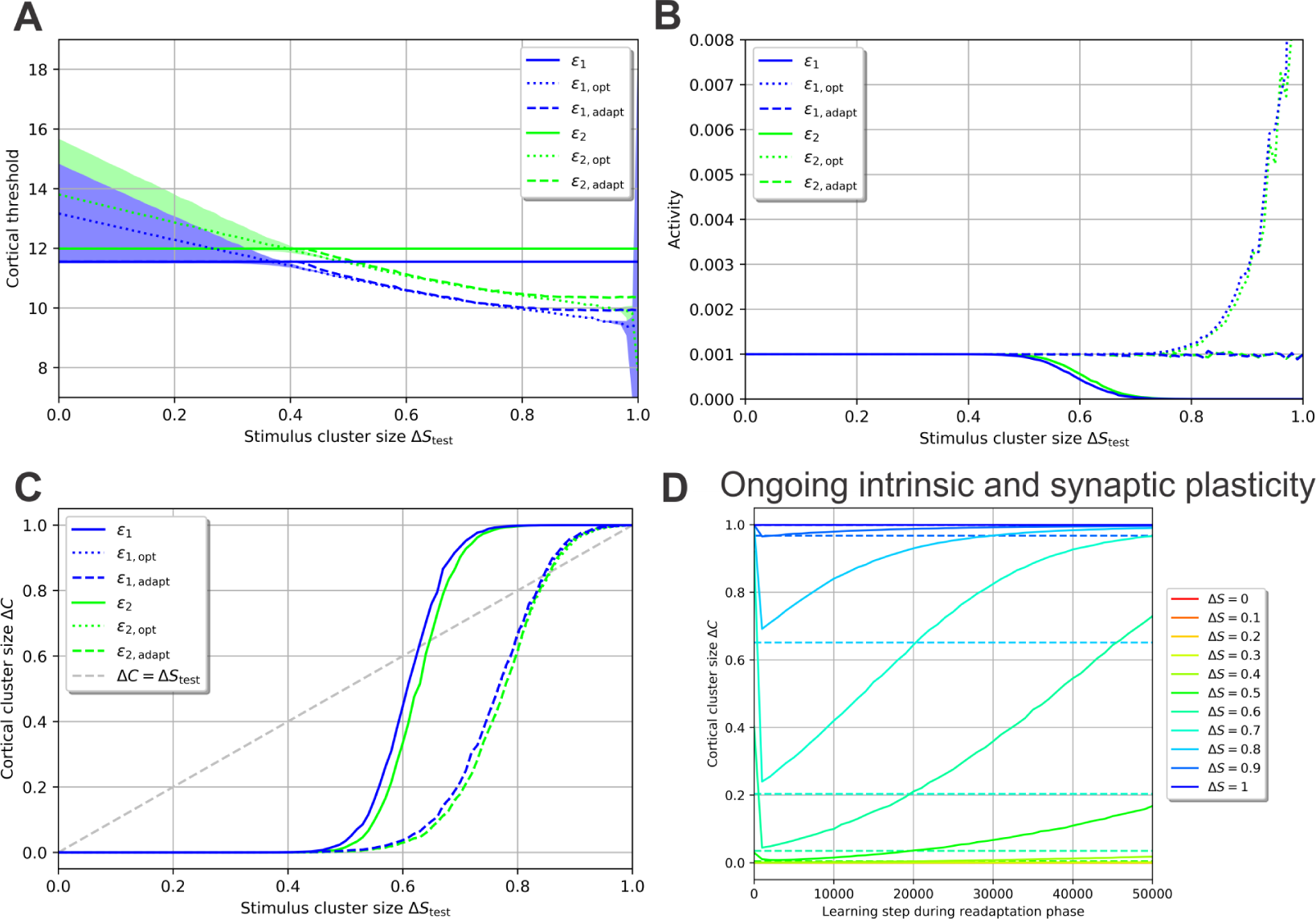
A second learning phase - the readaptation phase - enables the neuronal system to readapt to arbitrary noise levels using intrinsic plasticity. **(A-C)** After learning without noise, a second learning phase with the noise level Δ*S*_test_ and only intrinsic plasticity active enables the thresholds to readapt from the values after the first learning phase *ε*_*j*_ (solid lines) to adapted values *ε*_*j*,adapt_ (dashed lines) close to the optimal threshold values *ε*_*j*,opt_ (dotted lines) increasing performance. Blue: neuron 1; Green: neuron 2. (A) Δ*S*_test_-dependency of cortical thresholds; shaded areas indicate regimes of low error gradient (Fig 3C); (B) Δ*S* _test_-dependency of average activities; (C) Δ*S*_test_-dependency of total error. **(D)** If synaptic plasticity is present during the second learning phase as well, Δ*C* initially drops due to intrinsic plasticity and then increases with ongoing presentation of noisy stimuli indicating a disintegration of the synaptic structure. Dashed lines indicate Δ*C*-values for a second learning phase with intrinsic plasticity alone.

We therefore considered a second learning phase, the *readaptation phase*, which is conducted after the initial training or encoding phase is completed. In the readaptation phase, the stimulus cluster size will be the same that the performance is tested for, i.e. Δ*S*_test_. For now, synaptic plasticity is deactivated as we will only focus on intrinsic plasticity adapting the cortical firing thresholds. To implement this readaptation phase, after the first learning phase is completed, we repeatedly presented one noisy pattern 𝒮^*ν*^ per cluster using a stimulus cluster size Δ*S*_test_. Threshold adaptation was stopped when the mean of all cortical thresholds changed by less than 0.0001% in one step, which resulted in less than 7000 steps for each Δ*S*_test_. As expected, intrinsic plasticity adjusts the firing thresholds during this second phase so as to achieve the target firing rate ℱ^*T*^ for all Δ*S*_test_ (dashed lines; Fig 4B). Furthermore, the adapted thresholds are similar to the optimal thresholds (dashed lines; Fig 4A). This leads to a near-optimal classification performance, which is considerably better than without a readaptation phase (Fig 4C). Importantly, if synaptic plasticity is also present during this second learning phase, Δ*C* increases dramatically with ongoing readaptation (solid lines; Fig 4D). The initial drop of Δ*C* is due to intrinsic plasticity (dashed lines show final Δ*C*-values for intrinsic plasticity alone), while synaptic plasticity leads to a prolonged deterioration of the previously learned synaptic structure if stimuli are too noisy. We therefore conclude that the network has to maintain the synaptic weight structure during the readaptation phase, which we recreate by turning synaptic plasticity off. By doing so, the neuronal system can reliably adjust to stimuli of various noise levels using intrinsic plasticity for adapting the excitability of neurons.

### Plastic networks in noisy environments

Up to now, we have shown that a sparse, expansive network can learn the underlying organization of noise-free stimuli (Δ*S*_learn_ = 0) by means of synaptic and intrinsic plasticity. Afterwards, a readaptation phase with intrinsic plasticity alone enables the network to readapt to any arbitrary level of noise Δ*S*_test_ (Fig 4A-C). However, if synaptic plasticity is active during the readaptation phase, the noise of stimuli leads to a disintegration of the synaptic structure (Fig 4D). Therefore, it is unclear whether the network can also learn the organization of stimuli from noisy - instead of noise-free - stimuli by using synaptic plasticity.

To test this, we now investigate the effect of noisy stimuli during training in the encoding phase (i.e. Δ*S*_learn_ > 0). To do so, we present one noisy stimulus pattern 𝒮^*ν*^ per cluster in each learning step *L* using a stimulus cluster size Δ*S*_learn_. Remarkably, even for noise levels of Δ*S*_learn_ = 0.2 cortical neurons show neuronal and synaptic dynamics (Fig 5A,B) comparable to noise-free learning (Fig 2B,D). Synaptic weights and firing thresholds become correlated to the static, structured network (Fig 5E,F) to a similar degree (Fig 2C,E). Nevertheless, due to the noise of the stimuli, some cortical neurons do not manage to separate one stimulus cluster from all others (Fig 5D). Consequently, multiple clusters trigger the Hebbian term of synaptic plasticity (Eq 6) such that all synaptic weights approach a medium value (Fig 5C). These synaptic weights diminish the correlation to the static, structured synaptic weights as the final distribution is slightly broader (Fig 5E) than the one from learning without noise (Fig 2C). Furthermore, the cortical neurons without structured incoming synaptic weights (unimodal weight distribution) on average have a lower final firing threshold (blue outliers in Fig 5F).

**Fig 5.**
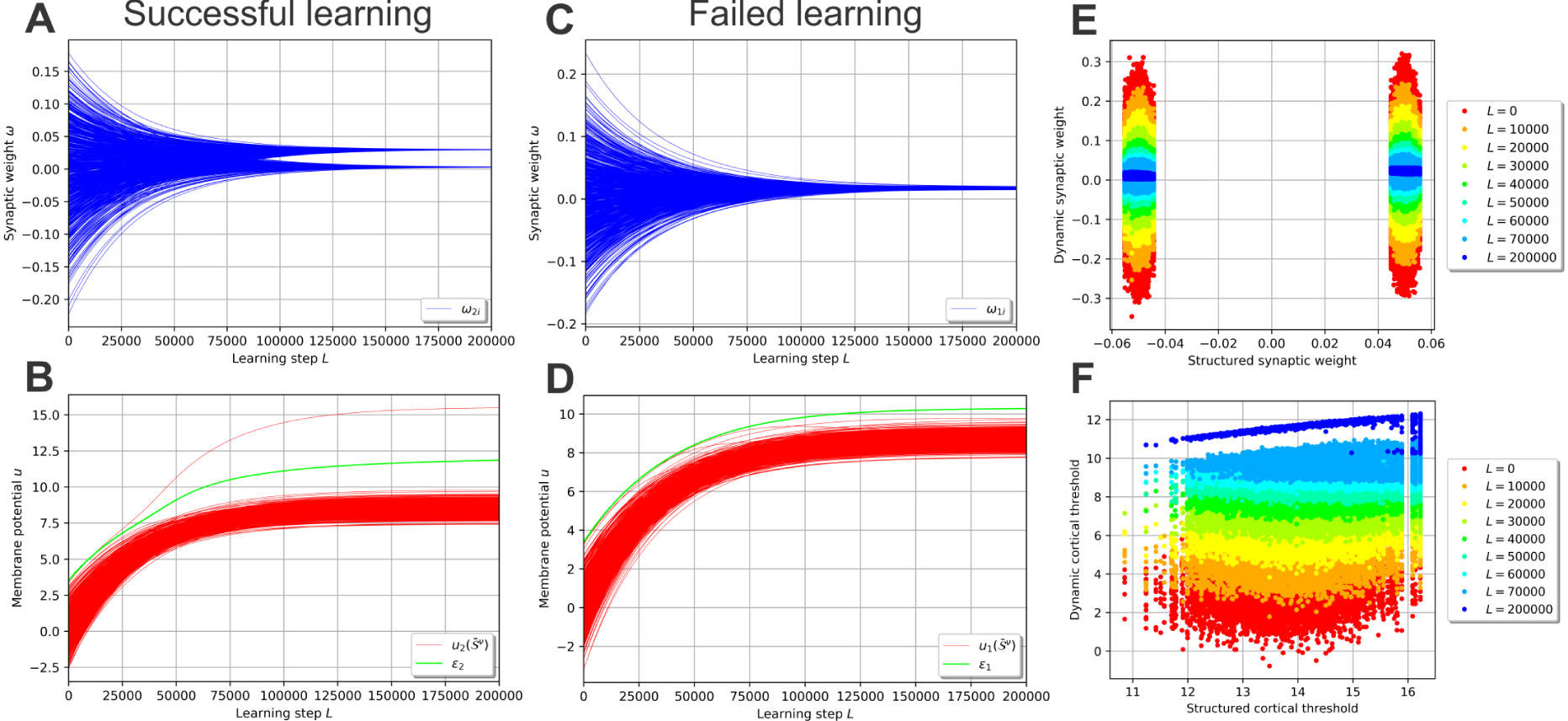
Self-Organization of the synaptic and neuronal structure in a noisy environment. The dynamics of synaptic and intrinsic plasticity enable the sparse, expansive network to learn the underlying organization of stimuli from noisy stimulus patterns (here Δ*S*_learn_ = 0.2). **(A,B)** The majority of cortical neurons develop a distribution of incoming synaptic weights (A) and membrane potential responses (B) similar to the ones learning without noise (Fig. 2B,D). Here shown for neuron 2. **(C,D)** However, the noise prevents some neurons to form a proper synaptic structure (C) yielding a firing threshold (D) which does not separate the membrane potential evoked by one cluster from the others. Therefore, these neurons are not tuned to one specific cluster. Here shown for neuron 1. **(E,F)** Overall, the network trained by noisy stimuli develops synaptic weights (E) and firing thresholds (F) similarly correlated to the static, structured network than the network trained without noise (Fig 2C,E). The few neurons that failed learning lead to a minor broadening of the distributions.

In general, low levels of noise (Δ*S*_learn_ ≲ 0.25) are tolerated by the network without large losses in performance (Fig 6A). The failed-learning cortical neurons (Fig 5C,D), which become more with higher noise levels, have a negative effect on the performance of the network. At Δ*S*_learn_ ≲ 0.25, the noise is so strong that the system is not able to recognize and learn the underlying organization of stimuli (that is, the existence of different clusters). Note that if the network learns from noisy stimuli, it can correctly classify stimuli which have a significantly higher level of noise (white area above orange dashed identity line). This result indicates that the network does not adapt specifically to only the noise level Δ*S*_learn_ it is learning from, but that the network generalizes across a broad variety of different noise levels Δ*S*_test_. For instance, although the network may learn from stimulus patterns with an average noise level of Δ*S*_learn_ = 0.1, it can reliably classify stimuli of noise levels Δ*S*_test_ from 0 to about 0.6 afterwards.

**Fig 6.**
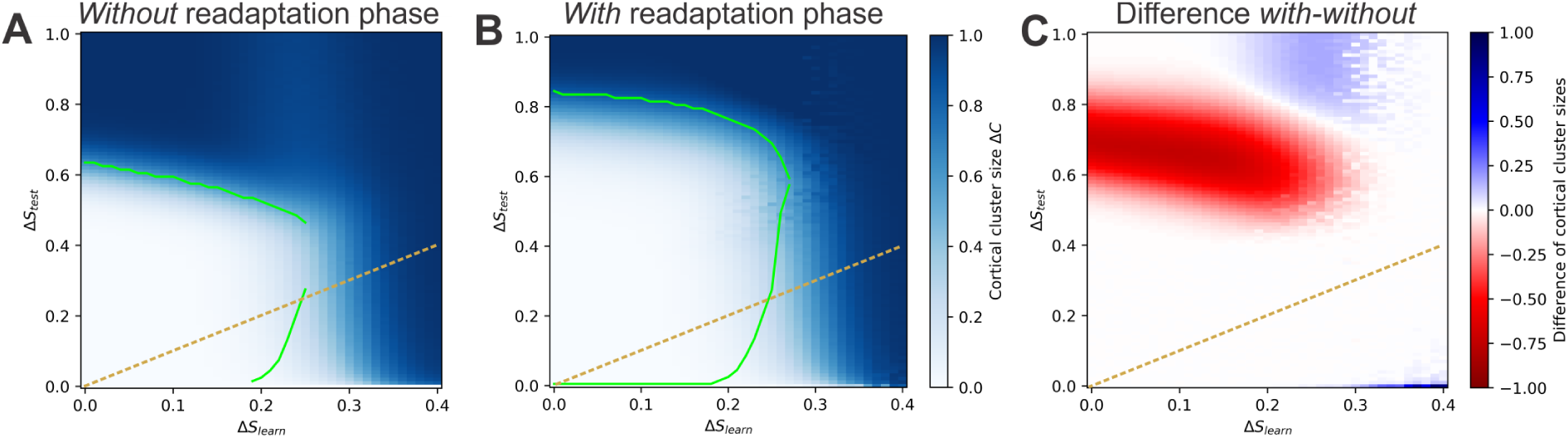
The network can reliably learn from noisy stimuli with and without a readaptation phase. **(A)** Despite the presence of noise Δ*S*_learn_ during learning, the network can learn the organization of stimuli and, after encoding, classify stimuli of even higher noise levels Δ*S*_test_. However, higher levels of Δ*S*_learn_ decrease the performance. **(B)** If the learning phase is followed by a readaptation phase using only intrinsic plasticity, the overall classification performance increases drastically. Now, stimuli with a noise level of up to Δ*S*_test_ ≈ 0.8 can be classified. **(C)** The readaptation phase leads to a large performance gain for medium and high noise levels Δ*S*_test_. Color code depicts the difference between the network without and with a readaptation phase. Red area represents a benefit by using the readaptation phase. (A-C) Orange, dashed line: identity line Δ*S*_learn_ = Δ*S*_test_

Furthermore, the performance of a network being successfully trained in a noisy environment can be drastically improved by a subsequent readaptation phase. Using this second phase in order to (re)adapt the neuronal excitabilities to the current level of noise Δ*S*_test_ enables the network to classify stimuli up to even higher noise levels of Δ*S*_test_ ≈ 0.8 (Fig 6B). Consequently, the readaptation phase provides a significant advantage for a large regime of stimulus cluster sizes (red area in Fig 6C). Even more so, stimulus clusters with sizes Δ*S*_test_ ∈ (0.6, 0.8) can only be classified by using the readaptation phase. The decrease in performance for noise levels between Δ*S*_learn_ ∈ (0.2, 0.3) and Δ*S*_test_ (0.8, 1.0) (blue area) is not crucial given the low level of performance (Fig 6A).

In summary, sparse, expansive networks can learn the clustered organization of noisy stimuli (underlying stimuli might be triangle, circle and cross like in Fig 7) by the interplay of synaptic and intrinsic plasticity in a self-organized manner. During the initial encoding phase, low levels of noise Δ*S*_learn_ can be tolerated by the system, while higher levels of noise obstruct the network’s ability to learn the organization of stimuli. After the encoding phase, the network can reliably classify noisy patterns of up to Δ*S*_test_ ≈ 0.6 if synaptic weights and neuronal firing thresholds are fixed 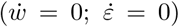. On the other hand, the performance decreases significantly if synaptic and intrinsic plasticity are allowed to modify the network’s structure during the presentation of these noisy stimuli 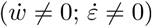. Interestingly, if the synaptic structure is maintained while the excitability of the cortical neurons can adapt 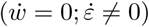, the network can successfully classify stimuli even in the presence of very high levels of noise (see Fig 7 bottom for examples). These results suggest that learning in the presence noise requires two distinct phases of plasticity: initial learning of the organization of environmental stimuli via synaptic and intrinsic plasticity in the encoding phase followed by the readaptation phase using only intrinsic plasticity in order to readapt to the current level of noise.

**Fig 7.**
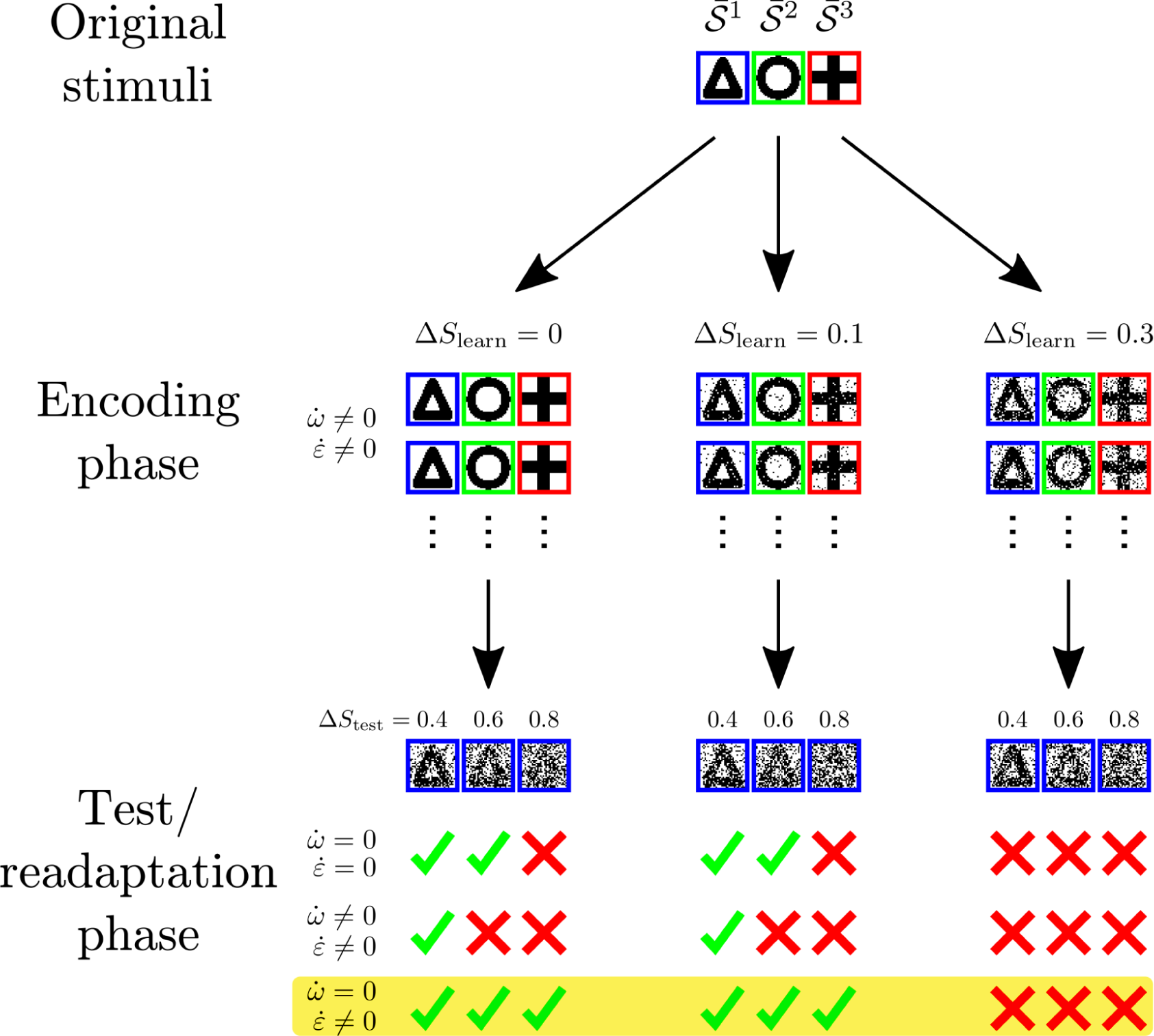
Schematic summary of results. Noisy patterns 𝒮^*ν*^ are repeatedly generated from underlying stimuli 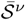 (e.g. a triangle, a circle and a cross) and imprinted on the stimulus layer (encoding phase). If the noise Δ*S*_learn_ is sufficiently small, synaptic and intrinsic plasticity lead to the formation of structure encoding the organization of stimuli (existence of different geometrical forms). After this initial learning phase, a second learning or readaptation phase enables the network to classify stimuli even in the presence of very high levels of noise Δ*S*_test_. Here, only intrinsic plasticity should be present 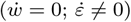. This suggests that learning is carried out in two phases: In the first phase, the *encoding phase*, synaptic weights develop to represent the basic organization of the environmental stimuli. This structuring of synaptic weights is most efficient if the noise Δ*S*_learn_ is low. In the second phase, the *readaptation phase*, learning is dominated by intrinsic plasticity while synaptic weights have to be maintained. The cortical firing thresholds are then able to quickly adapt to the current level of noise Δ*S*_test_. Thereby, intrinsic plasticity approximates the optimal thresholds for a given value of Δ*S*_test_ maximizing performance.

## Discussion

How do neuronal systems learn the underlying organization of the surrounding environment in realistic, noisy conditions? In this study, we have shown that sparse and expansive networks can reliably form the required neuronal and synaptic structure via the interplay of synaptic and intrinsic plasticity. Among others, our results indicate that the classification of diverse environmental stimuli in the presence of high levels of noise works best if the synaptic structure is more rigid than the neuronal structure, namely the excitabilities of the neurons.

The processes of synaptic consolidation could be a candidate to maintain the synaptic structure. A series of experimental and theoretical studies [40–44] show that only specific stimulation protocols trigger synaptic weight changes which are preserved for time scales of days (late-phase plasticity), while other changes decay after a few hours (early-phase plasticity). Therefore, neuronal activities during the encoding phase could trigger late-phase plasticity while recall stimuli during the test or readaptation phase cause mainly early-phase plasticity. To the best of our knowledge, neuronal excitabilities do not show any mechanisms of (long-term) consolidation and intrinsic plasticity can thus continuously adapt the neurons according to external stimuli. However, whether the processes of synaptic consolidation [41, 42, 44] yield the desired synaptic dynamics for the optimization of stimulus classification requires further investigations.

One of the major assumptions of this work, similar to a previous study [1], is that environmental stimuli are organized such that they can be grouped into clusters. Each of these clusters has the same size or noise level Δ*S*. Of course, in a natural environment each underlying stimulus can have a different level of noise and, therefore, each cluster *ν* could have a different size Δ*S*^*ν*^. However, if the synaptic structure has already been learned during the encoding phase, we expect that cluster-specific 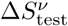 do not have an impact on the classification performance. This is because after initial learning each cortical neuron is selective to only one stimulus cluster (Fig 2D). In addition, only the noise level of this selected cluster defines the optimal firing threshold (Fig 3). Therefore, the firing threshold of each neuron can be tuned to its distinct, optimal threshold value, which is independent of the noise levels of other clusters. On the other hand, we expect that different 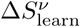 during the encoding phase will lead to over- and underrepresentations of stimulus clusters in the network. Since noise attenuates competition between clusters (Fig 5C,D), clusters with high 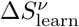 are less competitive and will subsequently be underrepresented. Nevertheless, the underrepresentation could be an advantage, as stimuli which are too noisy are less informative about the environment than others; consequently, the neuronal system attributes a smaller amount of resources (neurons and synapses) to them. However, the effect of cluster-specific noise on the neuronal and synaptic dynamics have to be investigated further.

Additionally, some stimulus clusters might be perceived more often than others. The corresponding representations would become larger than average, since their relevant synapses are strengthened more often by Hebbian synaptic plasticity, leading to a competitive advantage. Larger representations of more frequently perceived stimulus clusters might provide a behavioral advantage, as these clusters also need to be classified more often. However, the discrepancy between the frequency of such a cluster and the target firing rate of a cortical neuron responding to it might pose a problem. As intrinsic plasticity tries to maintain the target activity, the firing threshold would be placed so high that even slight noise could not be tolerated. One solution might be that neurons could have different target activities [45] and clusters are selected such that target activity and presentation frequency match. A different mechanism could be global inhibition. A single inhibitory neuron or population of neurons connected to all relevant cortical neurons could homeostatically regulate the activity of the cortical layer by providing inhibitory feedback. Such a mechanisms has been identified, for instance, in the Drosophila mushroom body [46, 47].

In this study, only one combination of three different plasticity rules was investigated. Of course, many more plasticity mechanisms are conceivable and have been widely studied [48–51]. One mechanism could be synaptic scaling regulating the synaptic weights instead of the neuronal excitability such that the neurons reach a certain target firing rate [24, 27, 52–54]. Possibly, synaptic scaling could be an alternative mechanism to intrinsic plasticity and the exponential decay term of synaptic plasticity, as scaling stabilizes synaptic dynamics, keeps the neuronal activities in a certain regime, and introduces competition.

It is usually assumed that homeostatic synaptic plasticity is required for competition [55, 56]. In the present study, however, competition arises from the interactions of Hebbian synaptic plasticity and homeostatic intrinsic plasticity alone. Homeostatic intrinsic plasticity maintains a certain activity of a given cortical neuron. Stimuli compete for this activity. If one stimulus gains an activity advantage, it will see synapses activated by it strengthened. This leads to less strengthening of other synapses, because the occurrence of Hebbian synaptic plasticity is limited by homeostatic intrinsic plasticity. Synapses will only subsequently be weakened due to homeostatic synaptic plasticity (exponential decay term), which does not interfere in the interaction between Hebbian synaptic and homeostatic intrinsic plasticity generating competition (S2 Fig). Consequently, the widely held opinion that homeostatic synaptic plasticity is required for competition might have to be revised.

Overall, we can answer the question of how networks learn to classify stimuli in noisy environments as follows: Learning takes place in two distinct phases. The first phase is the *encoding phase*. Hebbian synaptic and homeostatic intrinsic plasticity structure synaptic weights so as to represent the organization of stimuli, with each neuron becoming selectively responsive to a single stimulus cluster. Optimal synaptic structure is achieved if stimuli are noise-free. The second learning phase, called *readaptation phase*, ensues in an arbitrarily noisy environment. Here, synaptic weights have to be maintained in order to preserve the previously learned synaptic structure. Meanwhile, homeostatic intrinsic plasticity regulates the activity of neurons. The firing thresholds are thereby adapted to their optimal values, maximizing classification performance in the current environment (Fig 7).

## Methods

### Network and plasticity mechanisms

In this study, a two-layered feed-forward network of rate-based neurons is investigated (Fig 1A). The first layer, called stimulus layer, consists of *N*_*S*_ = 1000 neurons, while the second layer, called cortical layer, consists of *N*_*C*_ = 10000 neurons. Feed-forward synaptic connections exist from all stimulus to all cortical neurons. Their synaptic strengths are given by *ω*_*ji*_ where *j* = {1, …, *N*_*C*_} denotes the postsynaptic cortical neuron and *i* = {1, …, *N*_*S*_} the presynaptic stimulus neuron. No recurrent connections are present.

The neurons of the stimulus layer will act as input. As such, the firing rate 𝒮_*i*_ of stimulus neuron *i* will be set to either zero or one. Each input therefore is a pattern of firing rates 𝒮_*i*_ = {0, 1} on the stimulus layer. These firing rates elicit membrane potentials in the cortical neurons, which follow the leaky integrator equation 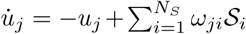. Transient dynamics are assumed to be negligible such that the membrane potential *u*_*j*_ of cortical neuron *j* can be simplified to

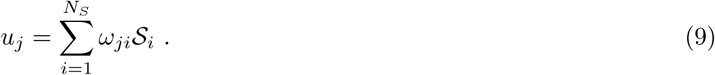

The membrane potential *u*_*j*_ will then be translated into a firing rate 𝒞_*j*_ of cortical neuron *j* via the sigmoidal transfer function

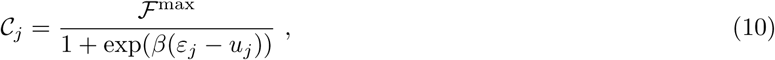

resulting in cortical firing rates between zero and ℱ^max^. The steepness of the sigmoidal function is given by *β* = 5, the maximum firing rate ℱ^max^ = 1, and the point of inflection *ε*_*j*_ is specific to each cortical neuron *j. ε*_*j*_ corresponds to a neuron-specific firing threshold determining the neuronal excitability.

Intrinsic plasticity regulates this neuron-specific firing threshold *ε*_*j*_. In order for each cortical neuron *j* to reach a target firing rate ℱ^*T*^ = 0.001, the point of inflection of the sigmoidal transfer curve follows the dynamics

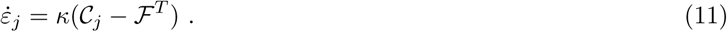

The parameter *κ* = 1 · 10^-2^ determines the adaptation speed of intrinsic plasticity. If the firing rate 𝒞_*j*_ of cortical neuron *j* is larger than the target firing rate ℱ^*T*^, the threshold *ε*_*j*_ increases such that 𝒞_*j*_ decreases (assuming the input stays constant), and vice versa.

The feed-forward synaptic connections *ω*_*ji*_ between the postsynaptic cortical neuron *j* and the presynaptic stimulus neuron *i* are controlled by unsupervised synaptic plasticity:

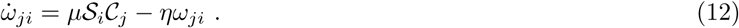

The parameters *μ* = 1 · 10^-5^ and *η* = 3 · 10^-8^ determine the speed of the Hebbian correlation learning term and the exponential decay of synaptic weights, respectively.

### Clustered stimuli

The structuring of the inputs and the analysis methods are similar to a previous work [1]. Here, sensory stimuli are grouped in *P* = 1000 clusters. Each cluster comprises different sensory impressions of the same environmental stimulus. Its main component is a characteristic neuronal firing pattern, called the central stimulus pattern 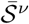, where *ν* = {1, …, *P*} denotes the cluster (Fig 1A,B). All central patterns are generated by assigning each stimulu neuron *i* for each stimulus cluster *ν* a firing rate 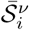 of either zero or one with equal probability. In addition to the central pattern, each cluster also contains noisy variants of the central pattern, called noisy patterns 𝒮^*ν*^. Noisy stimulus patterns are generated by randomly changing the central stimulus pattern’s firing rates from one to zero or vice versa with probability Δ*S/*2. Δ*S* thereby determines the level of noise and consequently the size of the stimulus clusters, and can range from zero (no noise) to one (no correlation remains). Furthermore, it is the normalized average Hemming distance of noisy stimulus patterns to their central stimulus pattern:

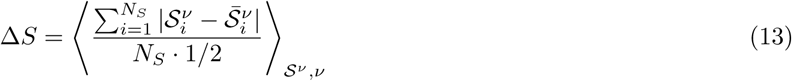

with the angular brackets denoting the average over all noisy stimulus patterns 𝒮^*ν*^ of all clusters *ν*.

All central and noisy stimulus patterns elicit central and noisy cortical patterns 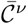 and 𝒞^*ν*^, respectively, in the cortical layer of the network. In analogy to Eq (13) the (uncorrected) size of the resulting cortical clusters can be defined as Δ*c* via

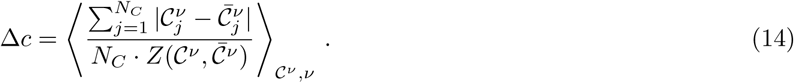

As the firing rates 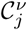 and 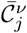 can take on values between 0 and 1, a more complex normalization *Z*(𝒞^*κ*^, 𝒞^*λ*^) for the patterns 𝒞^*κ*^ and 𝒞^*λ*^ is required:

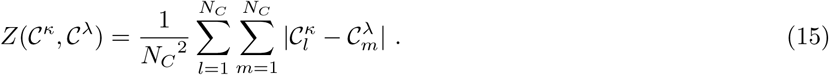

This normalization quantifies the average overlap two random cortical patterns with the same firing rates would have.

Being generated randomly, the central stimulus patterns are uncorrelated among each other. Due to the propagation of these patterns through the synaptic connections, however, the central cortical patterns might not be uncorrelated. In the context of noise reduction a more appropriate performance measure compensates for the introduced correlation. The cortical cluster size Δ*C* is therefore defined as:

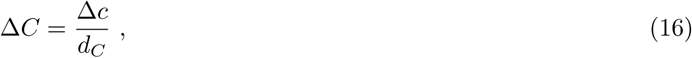

where the cortical cluster distance *d*_*C*_ is a measure of the correlation between central cortical patterns:

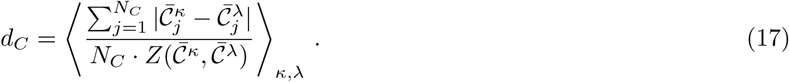

### Classification errors

In the following, the classification errors *e*_fp,*j*_ (false positives) and *e*_fn,*j*_ (false negatives) of single neurons will be set into relation with the cortical cluster size Δ*C*.

First, we will show that under certain assumptions the cortical cluster distance *d*_*C*_ = 1. The cortical cluster distance is defined as

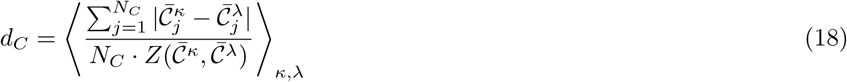

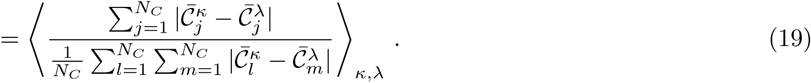

If we assume that cortical patterns are uncorrelated, the cortical cluster distance will be *d*_*C*_ = 1 by definition. We can demonstrate this by considering that every central stimulus pattern elicits a firing rate of 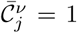 in ℱ^*T*^·*N*_*C*_ cortical neurons and 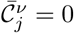 in all others, and every cortical neuron *j* fires with 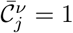 in response to ℱ^*T*^·*P* central patterns and with 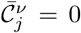 in response to all others. This is reasonably correct for successfully trained networks.Consequently:

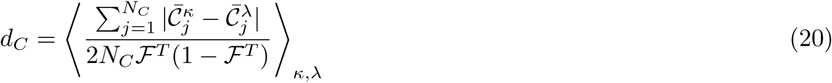

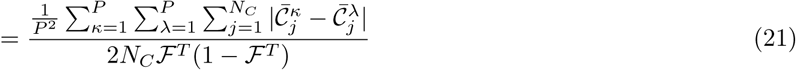

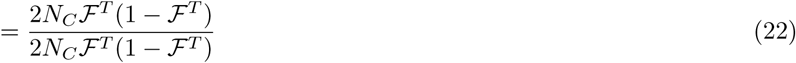

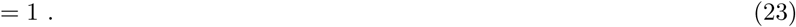

We therefore have:

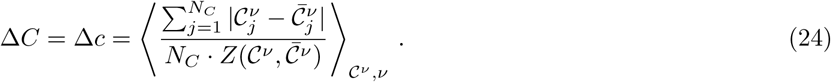

We will now demonstrate that under certain conditions the following relation holds:

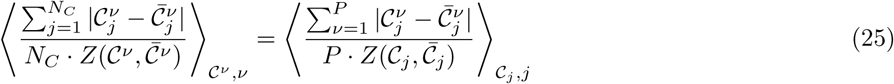

With

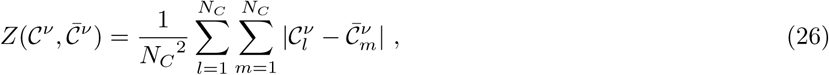

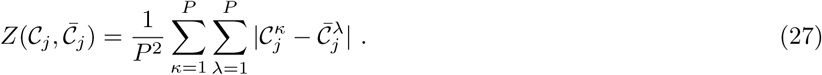

𝒞_*j*_and 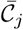 are vectors containing the firing rates of cortical neuron *j* in response to noisy/central patterns of all clusters. We will start from Eq 25 and extract a set of assumptions sufficient for the equation to hold: For a large system, i.e. *P, N*_*C*_ → ∞, the averages over 𝒞^*ν*^ and 𝒞_*j*_ can be dropped.

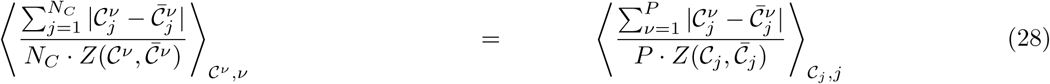

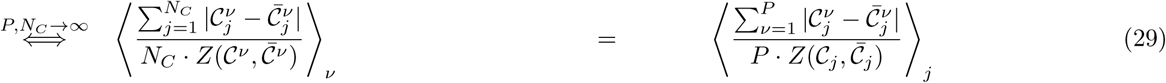

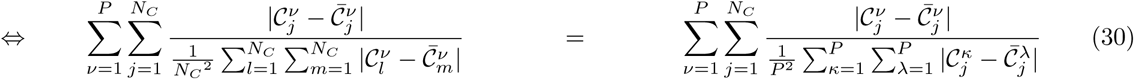

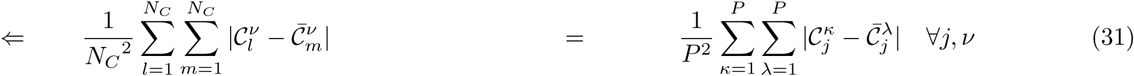

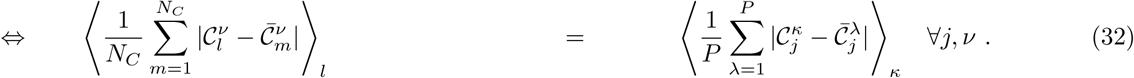

Again assuming every central pattern elicits a firing rate of 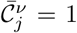 in ℱ^*T*^·*N*_*C*_ cortical neurons and 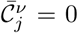 in all others, and every cortical neuron fires with 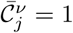 in response to ℱ^*T*^·*P* central patterns and with 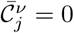 in response to all others, yields:

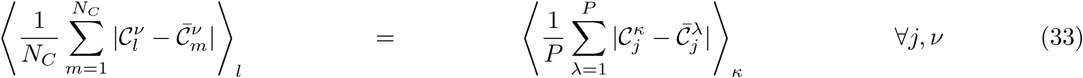

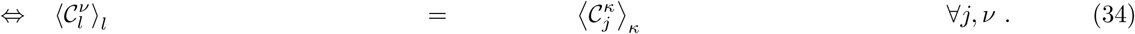

That is, for each Δ*S* every cortical pattern and every cortical neuron must have the same average firing rate.

Consequently, we now have:

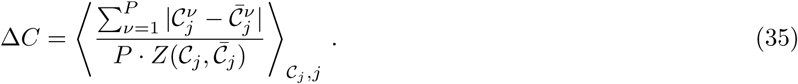

Further assuming that all clusters and all cortical neurons are comparable, the average over different neurons can be dropped:

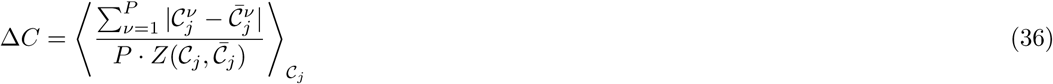

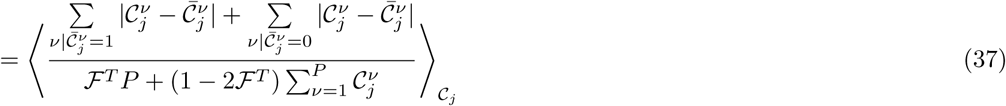

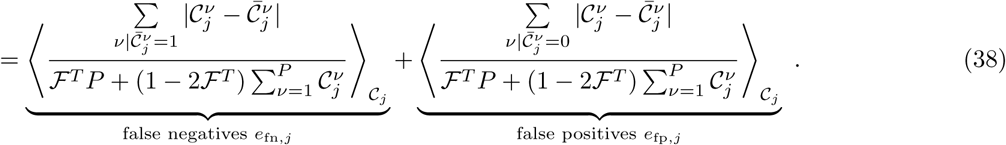

### Initialization

When initialised randomly, the synaptic weights *ω*_*ji*_ are drawn from a Gaussian distribution with mean 0 and variance 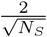.The synaptic weights can also be initialised in a structured manner according to 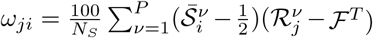 (similar to [1]). Here, ℛ^*ν*^ are cortical patterns that are generated using one of the following methods: For Fig 1 ℛ^*ν*^ are random patterns of ones and zeros where each pattern *ν* and each cortical neuron *j* has an activity of ℱ^*T*^. For all other results, ℛ^*ν*^ are computed via 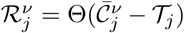, where Θ denotes the Heaviside function and the thresholds 𝒯_*j*_ are chosen such that each cortical neuron *j* achieves an activity of ℱ^*T*^. This results in cortical patterns ℛ^*ν*^ that are correlated to the central cortical patterns 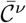 of an existing network.

The cortical membrane thresholds *ε*_*j*_ are then initialized such that each cortical neuron *j* achieves an average firing rate of the target firing rate ℱ^*T*^ at the central cortical patterns. In order to find the corresponding membrane thresholds *ε*_*j*_, the secant method is used with initial values of 0 and the mean of the highest and second highest (as ℱ^*T*^ *P* = 1) membrane potentials of cortical neuron *j*. If structured synaptic weights are used, this leads to *ε*_*j*_ close to the mean of the highest and second highest membrane potentials of neuron *j*.

### Implementation

Training is done by repeatedly presenting stimulus patterns for one time step Δ*t* = 1 each. One training step consists of parallel simulation of one stimulus pattern per cluster, ensuring that there is no chronological order of clusters. The stimulus patterns are generated using a stimulus cluster size Δ*S*_learn_ and the current learning step is denoted by *L*. The synaptic weights *ω*_*ji*_ and cortical thresholds *ε*_*j*_ are updated at the end of each training step according to 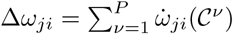 and 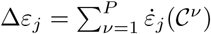 where *C*^*ν*^ denotes the cortical pattern of cluster 𝒞*ν* that was simulated in this learning step.

The cortical cluster size Δ*C* (cf. Eq 14 and Eq 16) in response to a stimulus cluster size Δ*S*_test_ is approximated using 10 noisy patterns per cluster.

If intrinsic plasticity is active during the testing phase, repeatedly, one noisy pattern per cluster is simulated using a stimulus cluster size Δ*S*_test_. After the mean of all cortical thresholds changed by less than 0.0001%, the cortical cluster size Δ*C* is calculated for the given stimulus cluster size Δ*S*_test_. In order to speed up its computation, we used the central cortical patterns 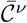 and cortical cluster distance *d*_*C*_ from ahead of the adaptation phase and were able to verify that this does not influence the results. The thresholds are reset to their previous values afterwards. The entire procedure is performed for all Δ*S*_test_, each requiring less than 7000 learning steps for the thresholds to converge.

## Supporting Information

**S1 Fig. Dependency of the classification error on the firing threshold for ΔS_test_** = **0**.**9.** At high levels of noise (here Δ*S*_test_ = 0.9), the actual value of the cortical firing threshold *ε* does not significantly influence the resulting classification error. This error remains close to one for all *ε*.

**S2 Fig. Competition without homeostatic synaptic plasticity.** The synaptic decay term was disabled during learning to demonstrate that competition arises from the interaction of Hebbian synaptic plasticity and homeostatic intrinsic plasticity alone. The exemplary neuron still becomes selective to only a single central pattern (left). All membrane potentials evoked by central pattern’s increase. However, the firing threshold (green) is also increased such that only one stimulus pattern results in a membrane potential above the threshold. Synapses activated by this pattern are increased, but as there is no reversive synaptic decay, they keep increasing (right). Note that this simulation was done with a ten-fold smaller network.

### Acknowledgments

This research was funded by the H2020-project Plan4Act (#732266) and by the German Funding Society’s (DFG) CRC 1286 (project C1).

